# Host cell cAMP-Epac pathway inhibition by hawthorn extract as a potential treatment for Chagas disease

**DOI:** 10.1101/2023.01.26.525677

**Authors:** Gabriel Ferri, Lucía R. Fernández, Guillermo Di Mario, Jorge A. Palermo, Martin M. Edreira

**Affiliations:** CONICET-Universidad de Buenos Aires, IQUIBICEN, Ciudad de Buenos Aires, Argentina; Laboratorio de Biología Molecular de Trypanosomas, Departamento de Química Biológica, Facultad de Ciencias Exactas y Naturales, Universidad de Buenos, Ciudad de Buenos Aires, Argentina; Universidad de Buenos Aires, Facultad de Ciencias Exactas y Naturales, Departamento de Química Orgánica, Buenos Aires, Argentina; CONICET-Universidad de Buenos Aires, Unidad de Microanálisis y Métodos Físicos Aplicados a la Química Orgánica (UMYMFOR), Buenos Aires, Argentina; Department of Pharmacology and Chemical Biology, School of Medicine, University of Pittsburgh, Pittsburgh, PA, USA

**Keywords:** Trypanosoma cruzi, cAMP signaling, Epac, Rap1b, Hawthorn, Crataegus, Vitexin, Host cell invasion

## Abstract

Benznidazole (BNZ) and nifurtimox (NFX), drugs used in the treatment of Chagas disease (CD), are effective in acute and congenital cases. However, due to the high toxicity of both drugs, the long duration of the treatment, the high doses, and the low effectiveness during the chronic phase, new therapies are needed. Recently, there has been an increase in alternative medicine and natural products popularity. Medicinal herbs emerge as a promising alternative for the development of new therapies against CD. The development of new active drugs requires the identification of new molecular targets. Host cell cAMP-Epac pathway plays a key role during *Trypanosoma cruzi* invasion. We have previously shown that Epac1 is required during the cAMP-mediated invasion of this parasite. Moreover, vitexin, a natural flavone that protects against ischemia-reperfusion damage, acts by inhibiting the expression of Epac and Rap1 proteins. Vitexin can be found in plants of the genus *Crataegus spp.*, traditionally known as hawthorn, that are of great interest considering their highly documented use as cardio-protectors. In this work, using HPLC-HRMS and MS2, we could confirm the presence of vitexin in an extract of *C. oxyacantha* (CO-EE). Interenstingly, treating cells with CO-EE, similar results for *T. cruzi* invasion than the ones observed for Epac1 specific inhibitor ESI-09 were observed. In addition, treated cells have a diminished activated Rap1b, suggesting that the extract could act through the cAMP-Epac signalling pathway. Most significantly, when using CO-EE in conjunction with NFX we observed an addition of the negative effects on the invasion, opening the possibility of decreasing the dosage/time currently used and thus alleviating the secondary side effects of available drugs, as well as the *per capita* treatment cost of CD.

## Introduction

Chagas disease (CD), also known as American trypanosomiasis, is a life-threatening illness caused by the flagellar protozoan *Trypanosoma cruzi*. Estimations indicate 7 million infected people and 75 million individuals at risk of contracting the disease, with an annual incidence of 30.000 cases and 10.000 deaths [1]. CD is endemic in 21 Latin American countries [2] and it belongs to the group of Neglected Tropical Diseases (NTD) since it mainly affects low-income populations, is one of the leading causes of mortality and chronic morbidity in developing countries and has a low economic allocation by research organizations [3]. Between 2009 and 2018, Chagas received only 0.67% of the total assigned to all NTD [4], which is approximately half of that received globally by other trypanosomiasis diseases, such as leishmaniasis or sleeping sickness [5]. Additionally, the human migration from endemic areas to Europe, North America, and the Western Pacific Region significantly increased the seroprevalence of *T. cruzi* infection outside Latin America [4]. Further, a worldwide financial cost of US$ 7.2 trillion per year is estimated [6] due to the loss of approximately 752,000 working days (premature deaths) and US$1-2 million in productivity [7].

CD develops in an acute and a chronic phase [5]. Once the *T. cruzi* trypomastigotes enter the body, acute CD occurs immediately. It is usually asymptomatic, with high parasite load circulating the blood [8]. Following the acute phase, most infected people enter a chronic phase and may remain asymptomatic for life without developing Chagas-related symptoms. Still, around 30–40% of infected people may present severe and sometimes life-threatening medical problems, namely chronic cardiac, digestive or mixed phase, and whose clinical manifestations will depend on the affected organ [9]. The cardiac form, also called Chagas heart disease, a dilated cardiomyopathy that involves a range of symptoms such as oedema, weakness and fatigue, palpitations, stroke and sudden death due to arrhythmias or progressive heart failure, is the most common manifestation among patients who develop symptoms at the chronic phase, representing the most cases of morbidity and mortality [10, 11].

There are two drugs currently in use for the treatment of CD, benznidazole (BNZ) and nifurtimox (NFX) [1, 12], that are effective in acute and congenital cases, or in the reactivation of the disease in immunosuppressed patients [13]. However, the treatment efficacy decreases with the progression of the disease [2]. Due to the high toxicity and low efficiency, especially during the chronic phase [14], the development of new suitable replacements is required. In addition, both compounds cannot be administered to pregnant women or people with kidney or liver failure, while NFX is also not recommended for patients with a history of neurological or psychiatric disorders [15].

The development of new active drugs against *T. cruzi* requires the identification of new molecular targets. Potential targets should be essential pathways in the parasite life cycle. Among these potential targets are signal transduction pathways, which could offer multiple components to be the target of new trypanocidal drugs [16]. Signaling pathways are both activated in the parasite [17] and during host cell invasion [18]. cAMP-mediated signalling plays a relevant role throughout the life cycle of *T. cruzi* [19] and could represent one of this potential molecular targets [20] since members of this pathway have been the target of numerous drugs before. For example, the modulation of adenylyl cyclases (AC) activity by agonists/antagonists targeting G protein coupled receptors (GPCRs) [21, 22] or forskolin derivatives drugs [23, 24], have been described in the treatment of heart failure. Diseases such as COPD, asthma, depression, schizophrenia, erectile dysfunction, psoriasis, and rheumatoid arthritis have been treated by the modulation of PDEs [25–29]. Further, Epac has recently been the target of different therapies through its specific inhibition, demonstrating a pharmacological effect in the prevention of invasion and metastasis of pancreatic and breast cancer and protection against fatal rickettsiosis [30].

In this regard, we have previously showed that Epac is the main mediator of cAMP-dependent invasion of the host cell [31] and, more recently, that this effect would require the activation and relocalization by the small GTPase of the Ras family, Rap1b (unpublished data).

As mentioned, the most frequent manifestation among patients who develop symptoms during the chronic phase is Chagas heart disease. This cardiac manifestation is considered an arrhythmogenic cardiomyopathy and is characterized by atrial and ventricular arrhythmias and a wide variety of conductive system abnormalities [10]. On the other hand, the presence of fibrosis and hypertrophy was observed in a 3D culture model of *T. cruzi* infected cardiomyocytes [32, 33] and, also, an increase in the frequency of tachyarrhythmias was recorded using infected VERO medium [34], suggesting that *T. cruzi* secreted proteins have the ability to potentiate arrhythmias, presumably by increasing Ca^2+^ intracellular levels. In turn, Epac-mediated activation is necessary for the stimulation of Ca^2+^ induced by Ca^2+^ release through ryanodine receptors (RyR) in cardiomyocytes [35], while Epac1 depletion protects against myocardial ischemia-reperfusion (I-R) damage, reducing infarct size and cell apoptosis [36]. Related to this, vitexin, a flavonoid compound derived from natural products that protects against I-R damage, may act by inhibiting the expression of Epac and Rap1 proteins, preventing mitochondria-mediated apoptosis [37]. Vitexin is present in a wide variety of herbs traditionally used in alternative medicine. In particular, plants of the Crataegus genus, commonly known as hawthorn, thornapple, May-tree, whitethorn, Mayflower or hawberry, are of great interest due to their highly documented use as cardioprotectors [38–42]. It has been shown that pre-treatment with a Crataegus tincture prevented the damage induced by isoproterenol in rat hearts [42]. It is interesting to note that isoproterenol-induced myocardial infarction damage is correlated with an elevation of the expression levels of Epac protein [43, 44], which could be done via vitexin or another compound of the hawthorn.

Considering heart as one of the most important targets in CD, and that the role play by the cAMP/Epac pathway in *T. cruzi* infection, we hypothesised that the inhibition of this pathway via natural cardio-protector compounds, such as vitexin, could be used as an alternative and/or complementary treatment against CD. Therefore, we performed a manual screening by analysing medicinal herbs databases and obtaining as a result herbs approved for human use that also protects against the damage produced by the parasite in the heart. Finally, we study these plants through extracts and infusions and determine their mechanism of action and their potential use as an alternative treatment for CD.

## Materials and Methods

### Cells and parasites

VERO (ATCC^®^ CCL-81^™^) and HELA (ATCC^®^ CCL-2^™^) cell lines were cultured in DMEM medium supplemented with Glutamax^TM^ (Gibco), 10% (v/v) FBS (Natocor), 100 U/ml penicillin and 0.1 mg/ml streptomycin (Sigma), and maintained at 37°C in a 5% CO_2_ atmosphere. Tissue culture-derived trypomastigotes forms (TCT) of *T. cruzi* Y strain were routinely maintained in VERO cells cultured in DMEM supplemented with 4% FBS and penicillin/streptomycin. Trypomastigotes were obtained from supernatants of infected VERO cells by centrifugation. First, the supernatant conditioned medium was centrifuged at low speed (500 g) to remove intact cells and cell debris Then, the supernatant obtained was centrifugated at 3,000 g for 15 min. and the pellet with the parasites was washed in PBS three times.

### Host cell transfection

A transient transfection protocol with polyethyleneimine (PEI) was used [45]. Briefly, cells were grown at about 60% confluence and incubated at 37 °C in a 5% CO_2_, 95% humidified air environment. Next day, cells were transfected with pCGN empty vector (EMPTY) or pCGN-HA-Rap1b (HA-Rap1) (kindly provided by Dr D. Altschuler, University of Pittsburgh, USA) using a ratio of 4:1 DNA:PEI mix in OptiMEM medium (Gibco). The mixture was kept for 30 min. at room temperature and then added to the cells and incubated at 37 C and 5% CO_2_. After 24h, cells were washed with PBS and complete medium (DMEM or Claycomb 10% FBS) was added. The transfected cells were used at 24h post-transfecction.

### Invasion assay

Cells were grown on glass cover slides in a 24 multi-well plate with DMEM 10% FBS for 24 hours at 2×10^4^ cells/well density at 37 °C, 5% CO_2_ and incubated with or without: 37,5 μM of the Epac1 specific inhibitor ESI-09 (Sigma), dilutions of hydroalcoholic extract of *C. oxyacantha* or 0.4% infusion of the aerial parts of *C. oxyacantha* plant. Cells were then washed and infected with trypomastigotes of the Y strain (moi 100:1) for 2 hours. Parasite were removed and cells incubated for 48 hs. Cells were fixed, stained with DAPI and infection level determined by fluorescence microscopy. Percentage of invasion and amastigotes/100 cells were calculated counting 3,000 cells, expressed as mean ± SD of three or more independent experiments and performed in triplicate. Infection of non-treated cells was considered as basal infection.

### GST Pull-down

HA-Rap1 transfected cells were incubated for 2 h with 300 μM of 8-Br-cAMP, 37,5 μM of ESI-09, 0.04% of CO-EE or respective controls. Detection of active Rap1 (GTP-bound) was performed through pull-down assays using a recombinant GST-RBD protein (GST fusion to the Rap1b-binding domain of the RalGDS protein, which only recognizes active Rap). A total of 1 mL bacteria lysates containing GST or GST-RBD were mixed by rotation with 40 μl 50% GSH-Sepharose at 4 °C for 1 h. The beads were centrifuged at 800 g for 2 min. at 4°C and washed with lysis buffer. Lysates from HA-Rap1 transfected cells pre-treated for 2h with 8Br-cAMP, infected with trypomastigotes of the Y strain (Tp Y) or mock infected (Ctrl) were incubated with RBD-glutathione-agarose resin for 1h at 4°C. Resin was washed and eluted with cracking buffer for WB analysis.

### Western Blot

After electrophoresis, the gel was equilibrated in 25 mM Trizma base, 192 mM L-1 glycine and 20% v/v methanol pH 8.3. Then, proteins were transferred to previously hydrated with methanol PVDF membranes (Amersham^™^ Hybond, GE Healthcare) in a vertical tank (Mini-PROTEAN^®^ Tetra Cell, Bio-Rad). After transfer, membranes were blocked with 20 mM L-1 Tris-HCl, 500 mM NaCl, 0.05% Tween and 5% non-fat milk, pH 7.5, incubated with anti-HA (Roche) antibody. After incubation, membrane was washed and incubated with rabbit horseradish peroxidase (HRP)-IgGs antibody (Santa Cruz Biotechnology), washed again and then revealed using 0.88 mg/ml luminol, 0.066 mg/ml p-coumaric acid, 6 mM H_2_O_2_, 100 mM Tris-HCl, pH 8.8 solution. Chemiluminescence was recorded with the C-DiGit scanner (LI-COR), and bands intensity were quantified with ImageJ software.

### HPLC-MS^2^

LC/MS analyses were performed on a RRLC Agilent 1200 using a Luna C18 column (3 μm, 2.0 × 100 mm; Phenomenex, Torrance, CA, USA). The mobile phase consisted of water containing 0.1% formic acid (A) and the solvent (B) was methanol. The flow rate was 0.3 mL/min, and the column temperature was set at 30 °C. Linear gradient elution was performed as follows: 10-75% B (0-25 min), 75-100 % B (25-26 min), 100% B (26-44 min). A diode array was used as detector coupled to a mass spectrometer. Triplicates of each sample were carried out.

Mass spectrometric analyses were performed using a Bruker MicrOTOF-Q II mass spectrometer (Bruker Daltonics, Billerica, MA, USA), equipped with an electrospray ion source. The instrument was operated at a capillary voltage of 4.5 kV with an endplate offset of 500 V, a dry temperature of 200 °C using N_2_ as dry gas at 6.0 L/min, and a nebulizer pressure of 3.0 bar. Multipoint mass calibration was carried out using a sodium formate solution from m/z 50 to 1200 in negative ion mode. Data acquisition and processing were carried out using the software Bruker Compass Data Analysis version 4.0 supplied with the instrument.

### Metabolites identification

Raw data was converted to ABF and pre-processed using the MS-DIAL freeware. ALL-GNPS library was used to search for matches between the features table obtained for the most intense ions and known metabolites. The results were manually cured comparing MS^2^ spectra of compounds already reported for the species.

### Docking

Blind Docking simulations of flavonoid compounds present in *C. oxyacantha* extract localized in Rap1b binding site of Epac. Structural alignment (cartoon and transparent electronic surface representation) between AF_Epac2, along with docked compounds, and crystal structure of Epac2-Rap1b complex (PDB ID: 4MGI). For better visualization of flavonoids, structure of AF_Epac2 was removed. Green: Epac2 regulatory region (CNBD + DEP); blue: Epac2 catalytic region (REM + RA + GEF domains); yellow: Rap1b.

## Results

There are reports in literature describing that vitexin may act by inhibiting the expression of Epac and Rap1 proteins [37, 46]. Vitexin is a flavone present in a wide variety of herbs traditionally used in alternative medicine. By analysing databases of Chinese medicinal herbs that can contain vitexin [47, 48] and crossing these data with herbs approved for human use [49, 50] that could also protect against the damage caused by *T. cruzi* in the heart [51], we found that *Crataegus spp*. plants, traditionally known as hawthorn or thornapple, were of great interest as a potential treatment against CD (Figure 1). HPLC-ESI(-)-HRMS and MS2 analysis confirmed the presence of vitexin in the extract of *C. oxyacantha* (CO-EE) commercially obtained (Droguería & Herboristería Argentina - Timos S.A.), as well as other flavonoids already reported for the species (Table 1) [52]. Remarkably, the pre-treatment of HELA cells with CO-EE produces a dose-dependent effect on *T. cruzi* invasion comparable to the Epac1 specific inhibitor (ESI-09) effect (Figure 2). Furthermore, cells pre-treated with an infusion made of aerial parts of *C. oxyacantha* plant (CO-Inf) showed a decrease in the invasion level similar to those obtained by pre-treating with CO-EE (Figure 3).

**Figure 1:**
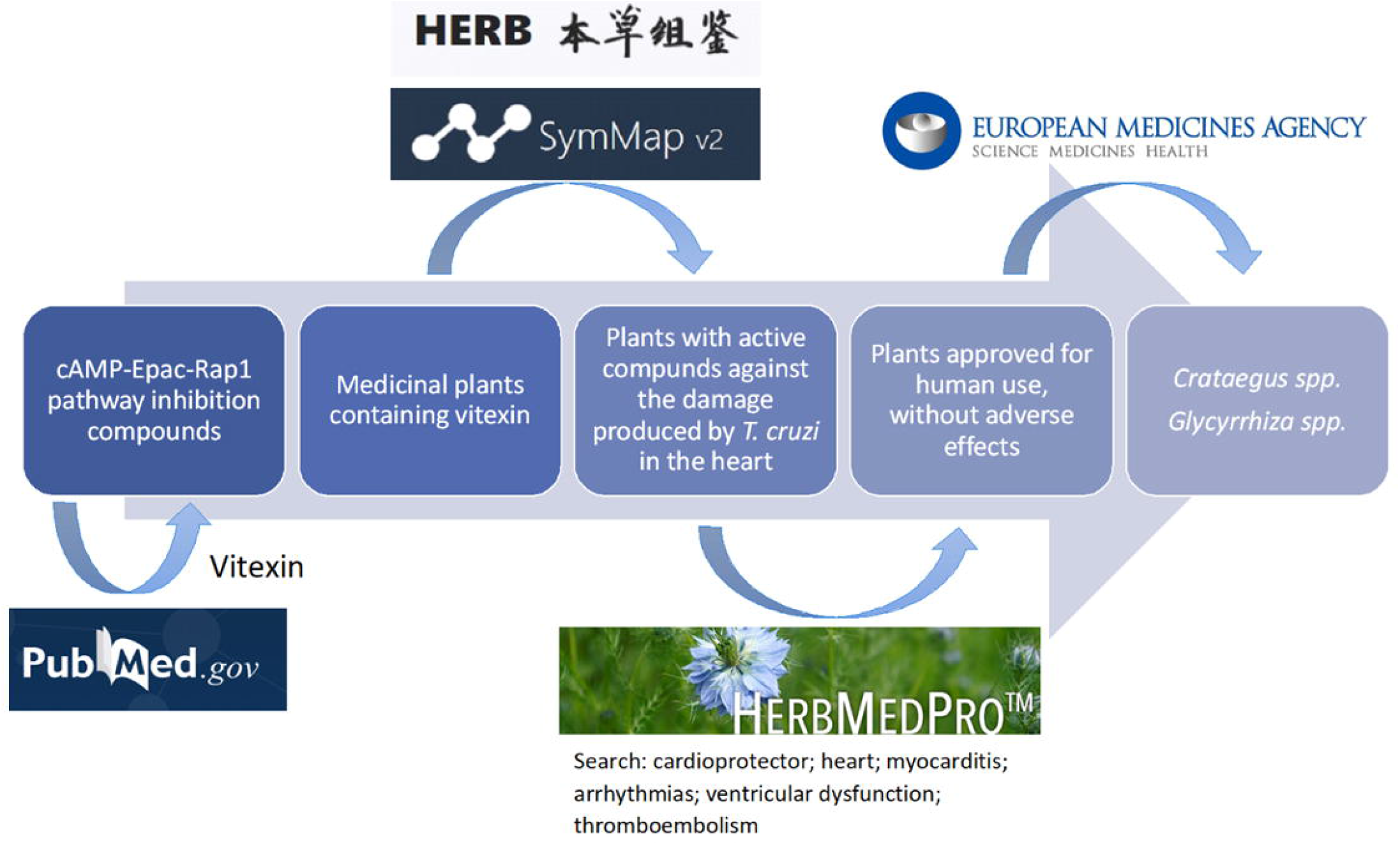
Manual screening in the search for medicinal herbs that can inhibit the cAMP-Epac-Rap1b pathway and protect against the damage caused by *T. cruzi* in the heart.

**Figure 2:**
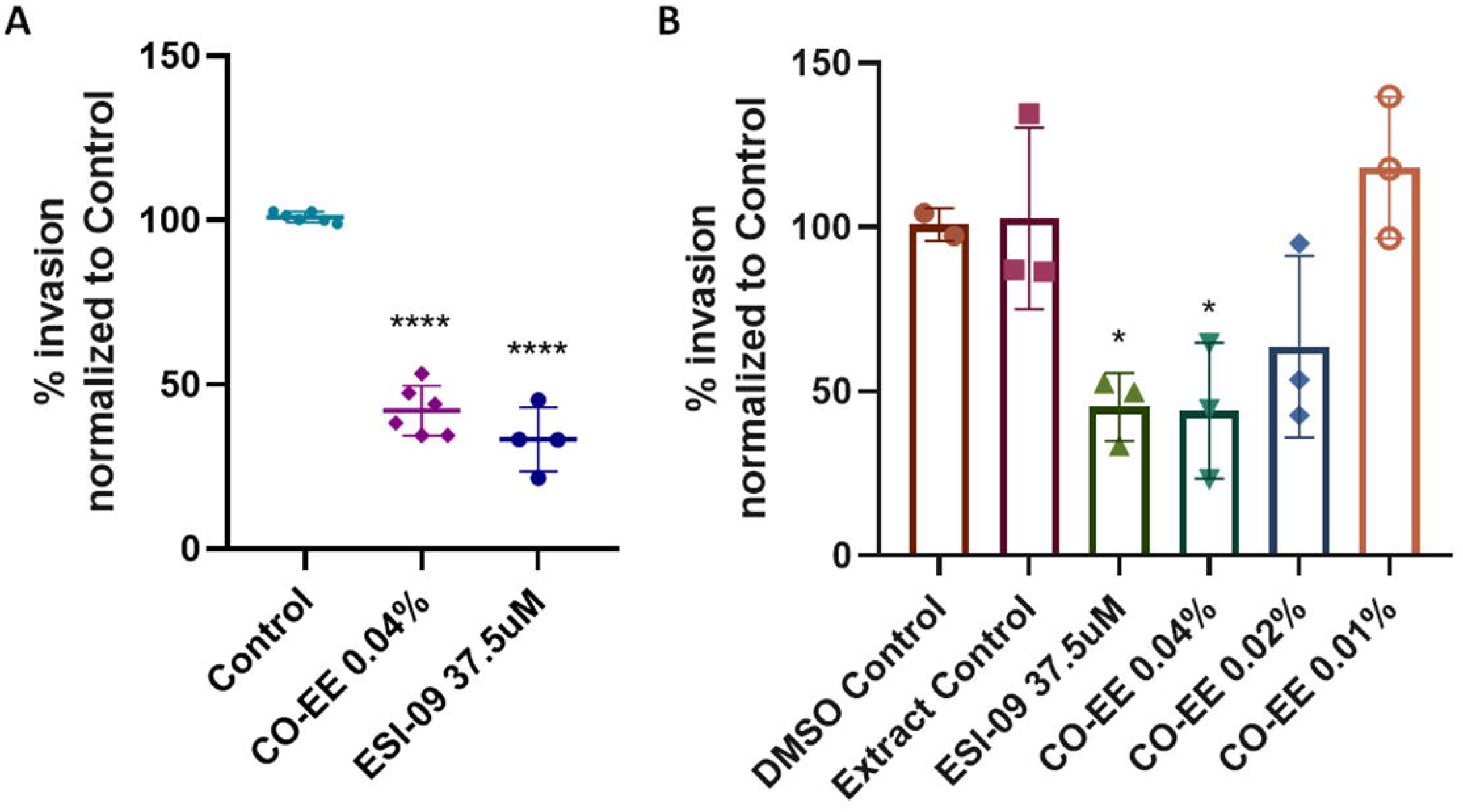
Pretreated HELA cells (1 h at 37.5 μM ESI-09 or different concentrations of CO-EE) were infected with trypomastigotes from *T. cruzi* Y strain (100:1 parasite to cell ratio for 2 h). 48 hs post-infection cells were fixed, stained with DAPI and percentage of invasion determined by fluorescence microscopy. Infection of untreated cells was considered as basal infection. Results are expressed as mean ± SD (n ≥ 3) **** p<0.0001, * p <0.1, ANOVA and Dunnett’s post-test. CO-EE: hydroalcoholic extract of *C. oxyacantha*.

**Figure 3:**
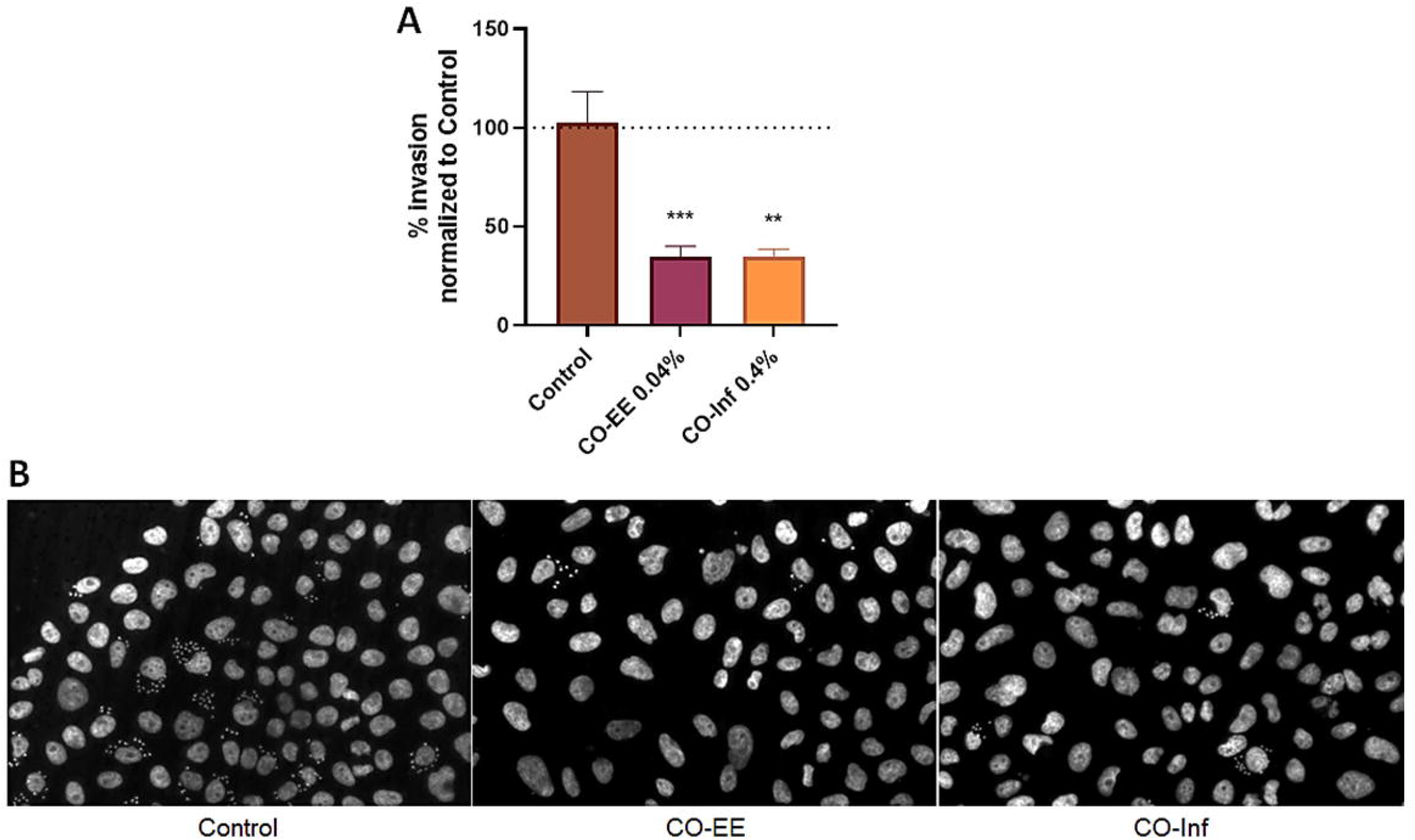
**A)** Pretreated HELA cells (1 h at 0.04% CO-EE or 0.4% CO-Inf) were infected with trypomastigotes from *T. cruzi* Y strain (100:1 parasite to cell ratio for 2 h). 48 hs post-infection cells were fixed, stained with DAPI and percentage of invasion determined by fluorescence microscopy. Infection of untreated cells was considered as basal infection. Results are expressed as mean ± SD (n ≥ 3) *** p<0.001, ** p <0.01, ANOVA and Dunnett’s post-test. **B)** Representative images of DAPI staining of infected cells pretreated with the indicated treatments. CO-Inf: infusion of the aerial parts of *C. oxyacantha* plant.

**Table 1.** Most intense [M-H]-ions observed in the extract of *C. oxyacantha* by HPLC-MS analysis with ESI ionization in negative mode. The identification was carried out by comparing the online libraries available in GNPS and subsequent manual curing considering the MS2 spectra and bibliography reported.

| Compound | tr<br>(min) | FM (M) | [M-H]<sup>-</sup> m/z<br>theoretical | [M-H]<sup>-</sup> m/z<br>observed | Error<br>(ppm) | MS<sup>2</sup> (% relative abundance) |
| --- | --- | --- | --- | --- | --- | --- |
| Quinic acid | 2.5 | C<sub>7</sub>H<sub>12</sub>O<sub>6</sub> | 191.0561 | 191.0560 | 0.5 | 1910(100), 173(12), 171(16), 153(10), 127(14) |
| Orientin | 15.8 | C<sub>21</sub>H<sub>20</sub>O<sub>11</sub> | 447.0933 | 447.0963 | -6.7 | 327(100), 357(68), 333(29), 297(16), 285(15) |
| Isoorientin | 16.1 | C<sub>21</sub>H<sub>20</sub>O<sub>11</sub> | 447.0933 | 447.0927 | 1.3 | 357(100), 327(72), 363(65), 297(15), 339(13), 285(13), 449(12) |
| Not identified | 16.2 | - | - | 593.1479 | - | 593 (100), 473(31), 327(7), 429(5) |
| Vitexin | 16.6 | C<sub>21</sub>H<sub>20</sub>O<sub>10</sub> | 431.0984 | 431.0954 | 6.9 | 311(100), 283(18), 341(14) |
| Vitexin 2''-O-rhamnoside | 16.9 | C<sub>27</sub>H<sub>30</sub>O<sub>14</sub> | 577.1563 | 577.1558 | 0.8 | 413(100), 577(34), 293(31), 457(10), 311(8) |
| Isovitexin | 17.3 | C<sub>21</sub>H<sub>20</sub>O<sub>10</sub> | 431.0984 | 431.0984 | 0 | 311(100), 341(52), 283(9), 295(9), 323(9), 313(7), 431(6), 353(5) |
| Isovitexin 2''-O-rhamnoside | 17.5 | C<sub>27</sub>H<sub>30</sub>O<sub>14</sub> | 577.1563 | 577.1593 | -5.1 | 577(100), 413(61), 293(35), 457(13), 323(5) |
| Quercetin 3-O hexoside | 18.0 | C<sub>21</sub>H<sub>20</sub>O<sub>12</sub> | 463.0882 | 463.0826 | 12.0 | 300(100), 301(34), 463(12) |
| Quercetin 3-O hexoside<br>(isomer) | 18.2 | C<sub>21</sub>H<sub>20</sub>O<sub>12</sub> | 463.0882 | 463.0836 | 9.9 | 300(100), 301(42), 463(10) |
| Isorhamnetin O-hexoside | 19.1 | C<sub>22</sub>H<sub>22</sub>O<sub>12</sub> | 477.1038 | 477.1046 | -1.7 | 314(100), 315(77), 300(64), 299(50), 477(41) |
| Quercetin | 21.5 | C<sub>15</sub>H<sub>10</sub>O<sub>7</sub> | 301.0354 | 301.0411 | -18.9 | Not fragmented |
| Isorhamnetin | 23.3 | C<sub>16</sub>H<sub>12</sub>O<sub>7</sub> | 315.0510 | 315.0438 | 22.8 | 300(100), 166(11), 227(9), 202(7), 245(7), 138(7) |

Regarding the mechanism of action, the inhibition of ESI-09 induced a similar decrease in invasion to CO-EE and no additive or synergistic effects were observed when incubating with the Epac inhibitor and with the *C. oxyacantha* extract simultaneously, suggesting that both compounds share the same mechanism of action (Figure 4). In line with this result, a lower level of activated Rap1b was detected in lysates from cells incubated with CO-EE, strongly suggesting that CO-EE would act through the Epac/Rap1b signaling pathway (Figure 5). Interestingly, not only vitexin, but several of the compounds found in the extract could bind to the Epac/Rap1b interphase (Figure 6) with a concomitant negative effect on the interaction and activation of Rap1b.

**Figure 4:**
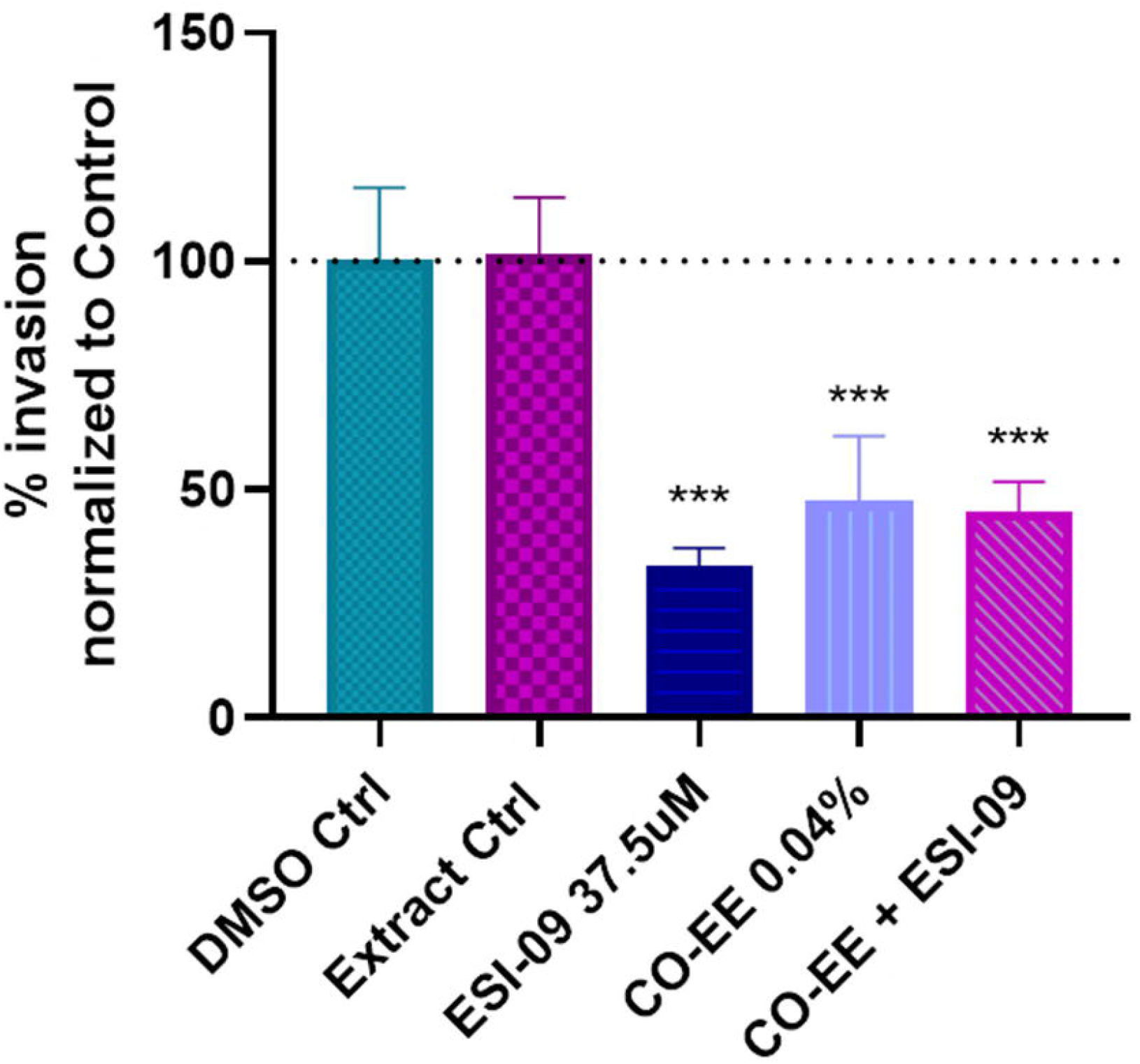
Pretreated HELA cells (1 h at 37.5 μM ESI-09, 0.04% CO-EE or both) were infected with trypomastigotes from *T. cruzi* Y strain (100:1 parasite to cell ratio for 2 h). 48 hs post-infection cells were fixed, stained with DAPI and percentage of invasion determined by fluorescence microscopy. Infection of untreated cells was considered as basal infection. Results are expressed as mean ± SD (n ≥ 3) *** p<0.001, ANOVA and Dunnett’s post-test.

**Figure 5:**
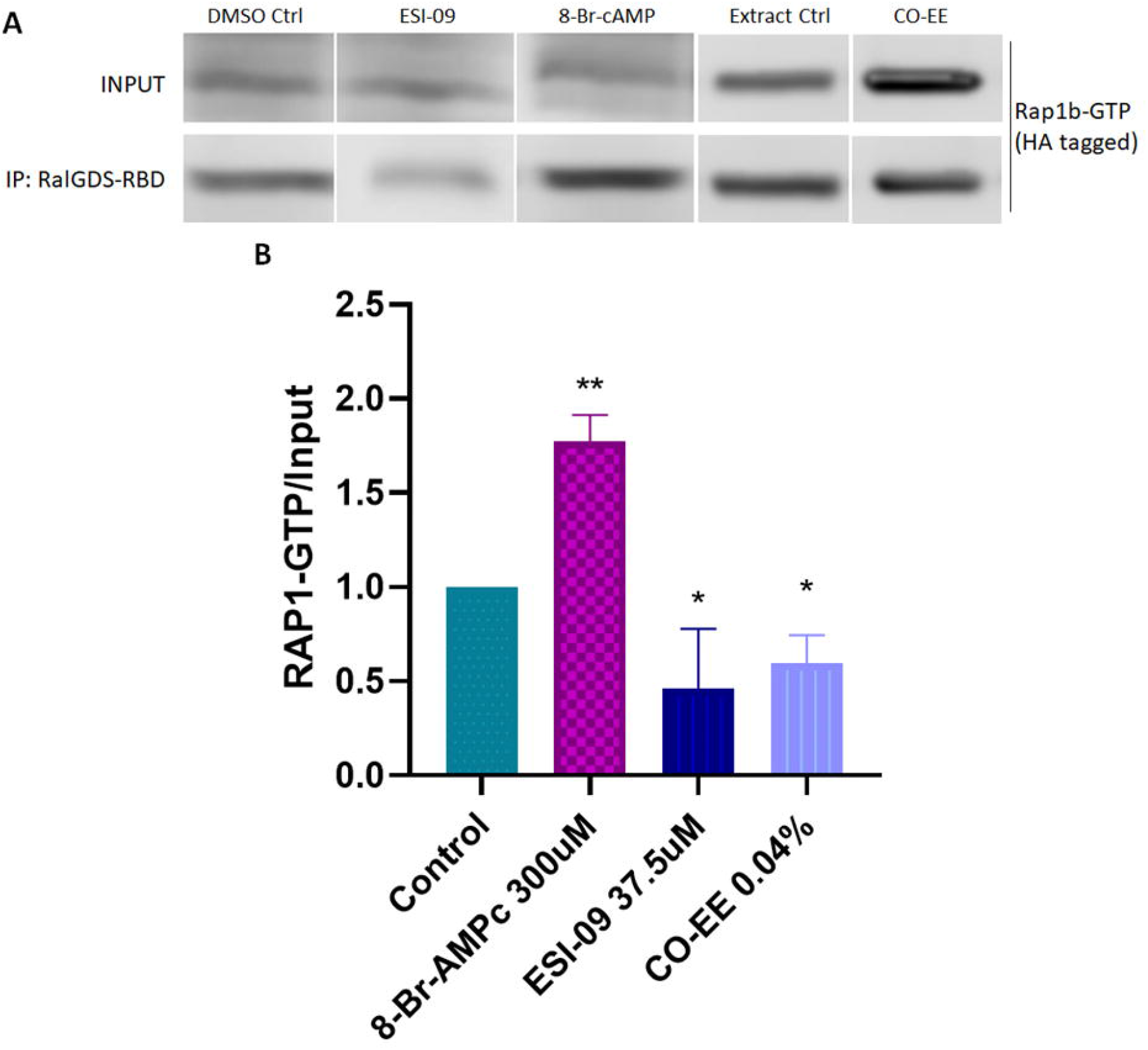
Rap1b pull-down assays. **A)** HA-Rap1 transfected HELA cells were incubated for 2 h with 8-Br-cAMP 300 μM, ESI-09 37.5 μM, 0.04% of CO-EE or control. Then, cells were lysed and pull-down assay with glutathione-agarose resin performed for 1 h at 4°C. Resin was washed and eluted with cracking buffer for WB analysis. **B)** Bands were quantified and normalized against the input using ImageJ cell software. Results are expressed as mean ± SD (n≥3) * p<0.05, t student test.

**Figure 6:**
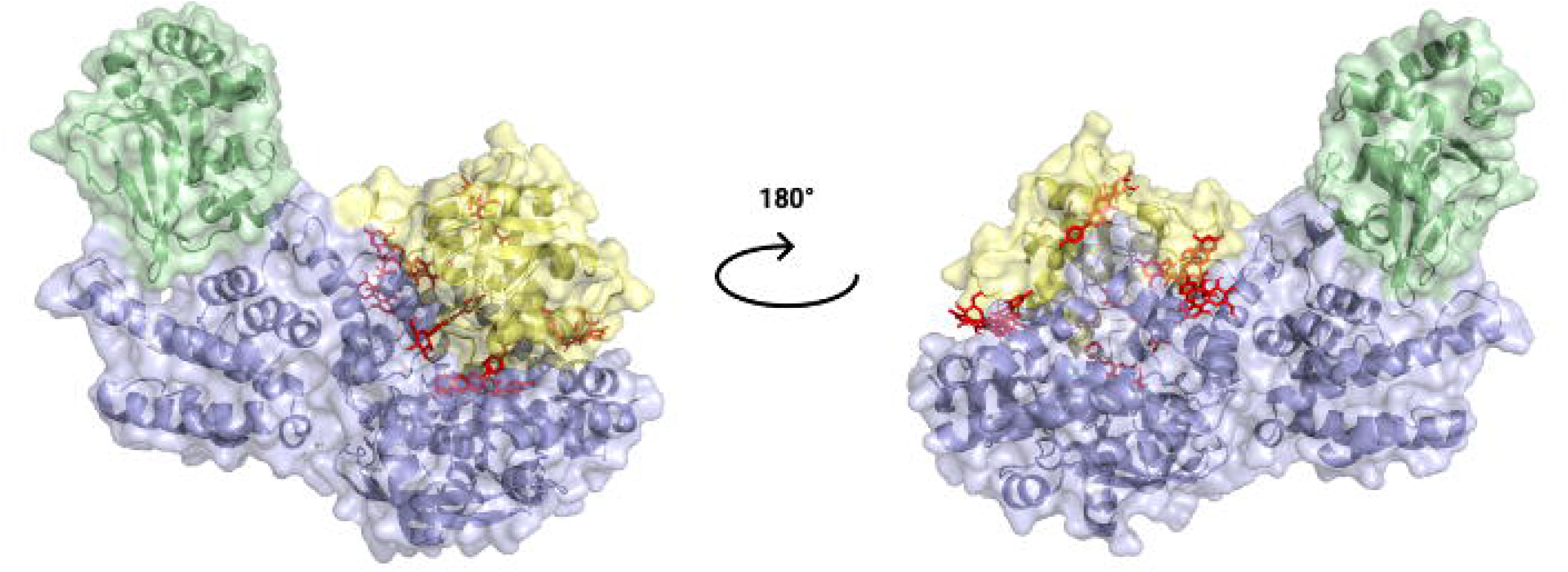
Blind Docking simulations of flavonoid compounds present in *C. oxyacantha* extract localized in Rap1b binding site of Epac. Structural alignment (cartoon and transparent electronic surface representation) between AF_Epac2, along with docked compounds, and crystal structure of Epac2-Rap1b complex (PDB ID: 4MGI). For better visualization of flavonoids, structure of AF_Epac2 was removed. Green: Epac2 regulatory region (CNBD + DEP); blue: Epac2 catalytic region (REM + RA + GEF domains); yellow: Rap1b.

Finally, *in vitro* invasion assays in cells pre-treated with CO-EE and then incubated for 48 hours with or without NFX showed that both compounds could act by different pathways since additive effects on the invasion level were detected (Figure 7), opening the possibility of a co-treatment of CD.

**Figure 7:**
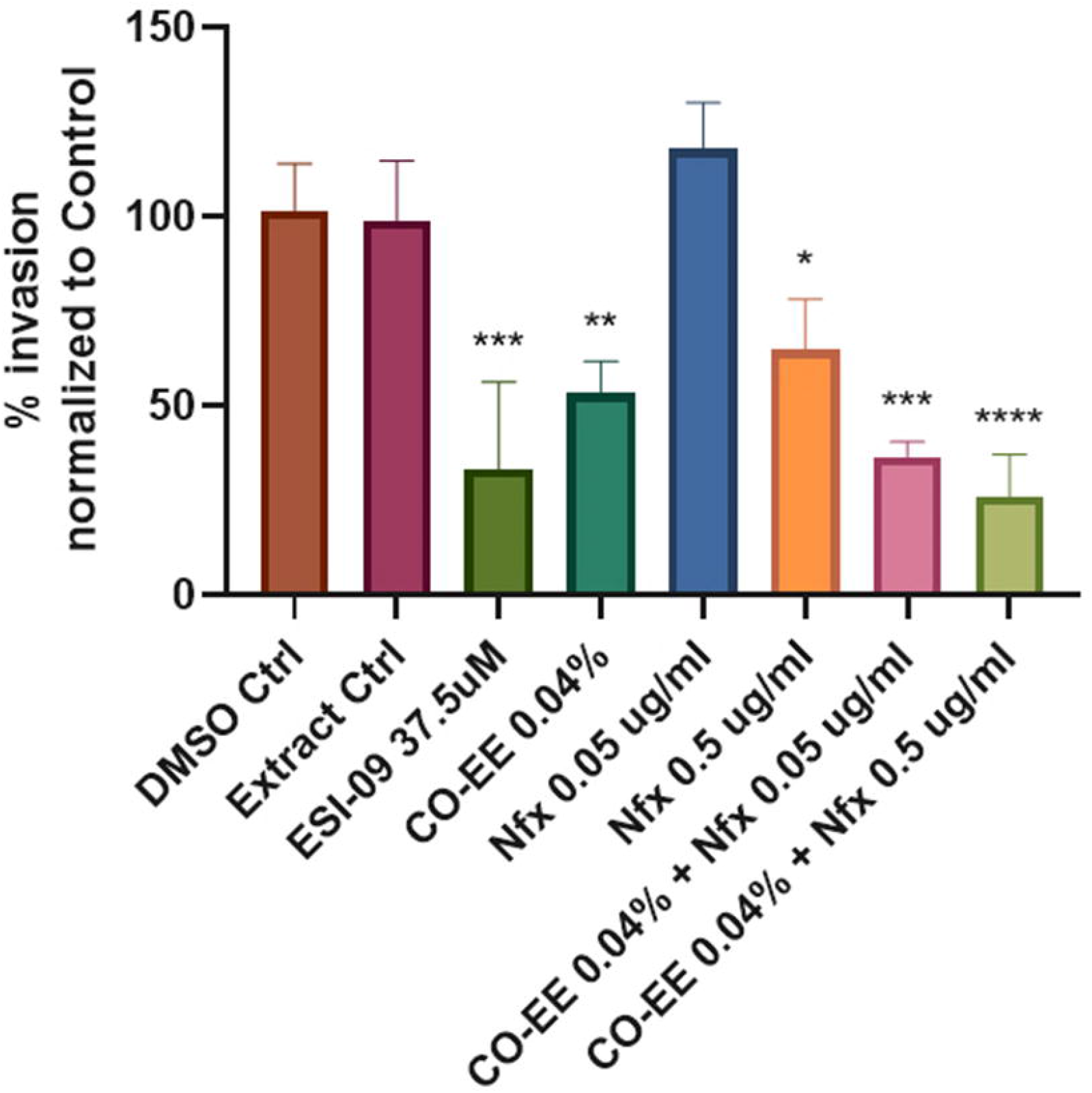
Pretreated HELA cells (1 h at 37.5 μM ESI-09 or 0.04% CO-EE) were infected with trypomastigotes from *T. cruzi* Y strain (100:1 parasite to cell ratio for 2 h). The medium was change to DMEM medium with different concentrations of NFX or DMSO as control and incubated 48 hs. Then cells were washed, fixed, stained with DAPI and percentage of invasion determined by fluorescence microscopy. Infection of untreated cells was considered as basal infection. Results are expressed as mean ± SD (n ≥ 3) *** p<0.001, ** p <0.01, * p <0.1, ANOVA and Dunnett’s post-test.

## Discussion

The inhibition of the cAMP/Epac/Rap1b pathway as a therapeutic target for CD was evaluated. During the chronic phase the most frequent manifestation is Chagas heart disease. *T. cruzi* secreted proteins can potentiate arrhythmias, presumably by increasing Ca^2+^ intracellular levels. In fact, in cardiomyocytes, Epac-mediated activation is necessary for the stimulation of Ca^2+^ induced by Ca^2+^ release through RyR [35], while Epac1 depletion protects against myocardial I-R damage [36]. Related to this, vitexin protects against I-R damage and may act by inhibiting the expression of Epac and Rap1 proteins [37]. Here, we showed that the hydroalcoholic extract of *C. oxyacantha* (CO-EE) provides protection against the invasion of *T. cruzi* in HELA cells *in vitro*.

We found that the pre-treatment of the cells with CO-EE produced a decrease in *T. cruzi* invasion. In addition, no additive effects were seen when using CO-EE and ESI-01, suggesting that both treatments share the same mechanism of action. Accordingly, CO-EE treated cells showed a decrease in Rap1b activation, confirming that the extract would inhibit the cAMP/Epac/Rap1b pathway. Several compounds present in the extract might be mediating this negative modulation on Rap1b, potentially by inhibiting Epac/Rap1b interaction.

On the other hand, when using CO-EE together with NFX, we observed an effect addition on the invasion, confirming that both drugs could act on different targets and opening the possibility of reducing dosage/time of NFX, which would contribute to avoiding adverse effects, as well as reducing the cost *per capita* of the treatment. The fact that CO-EE is used to treat cardiac pathologies through its antioxidant and antiarrhythmic properties [42, 53, 54] grants this medicinal herb unique characteristics in the potential treatment of CD, being able to prevent the invasion and spread of *T. cruzi* in the tissue while treating the damage caused by this parasite in the heart. Deciphering the cAMP signaling mechanisms activated by *T. cruzi* will provide a deeper insight into the mechanisms of host cell invasion mediated by this cyclic nucleotide. Considering that there is a wide variety of therapies that target cAMP-mediated signaling [55], revealing the actors of the cAMP/Epac1/Rap1b pathway could provide an attractive set of new therapeutic targets for the repositioning or development of antiparasitic drugs against CD.

## Author Contributions

GF, LRF and MME conceived, planned, and designed experiments. GF and LRF conducted experiments. GF and MME analysed the data and wrote the manuscript. All authors contributed to the article and approved the submitted version.

## Data availability

Data sharing not applicable to this article as no datasets were generated or analysed during the current study.

## References

1. WHO (2020) Ending the neglect to attain the Sustainable Development Goals: A road map for neglected tropical diseases 2021–2030, Control of. World Health Organization - Licence: CC BY-NC-SA 3.0 IGO

2. Pérez-Molina JA, Molina I (2018) Chagas Disease. Lancet. https://doi.org/10.1016/S0140-6736(17)31612-4

3. Bonney KM (2014) Chagas disease in the 21st Century: A public health success or an emerging threat? Parasite 21:. https://doi.org/10.1051/parasite/2014012

4. Sangenito LS, Branquinha MH, Santos ALS (2020) Funding for Chagas Disease: A 10-Year (2009-2018) Survey. Trop Med Infect Dis 5:. https://doi.org/10.3390/tropicalmed5020088

5. Echeverría LE, Marcus R, Novick G, et al (2020) WHF IASC Roadmap on Chagas Disease. Glob Heart 15:26. https://doi.org/https://doi.org/10.5334/gh.484

6. Lee BY, Bacon KM, Bottazzi ME, Hotez PJ (2013) Global economic burden of Chagas disease: a computational simulation model. Lancet Infect Dis 13:342–348. https://doi.org/10.1016/S1473-3099(13)70002-1

7. Conteh L, Engels T, Molyneux DH (2010) Socioeconomic aspects of neglected tropical diseases. Lancet 375:239–247. https://doi.org/10.1016/S0140-6736(09)61422-7

8. Menezes C, Costa GC, Gollob KJ, Dutra WO (2011) Clinical aspects of Chagas disease and implications for novel therapies. Drug Dev Res 72:471–479. https://doi.org/10.1002/ddr.20454

9. Matsuda NM, Miller SM, Evora PRB (2009) The chronic gastrointestinal manifestations of Chagas disease. Clinics 64:1219–1224. https://doi.org/10.1590/S1807-59322009001200013

10. Nunes MCP, Beaton A, Acquatella H, et al (2018) Chagas Cardiomyopathy: An Update of Current Clinical Knowledge and Management: A Scientific Statement From the American Heart Association

11. Martinez F, Perna E, Perrone S V, Liprandi AS (2019) Chagas Disease and Heart Failure: An Expanding Issue Worldwide. Eur Cardiol Rev 14:82–88. https://doi.org/10.15420/ecr.2018.30.2

12. Rodriques Coura J, de Castro SL (2002) A critical review on Chagas disease chemotherapy. Mem Inst Oswaldo Cruz 97:3–24. https://doi.org/10.1590/s0074-02762002000100001

13. Lidani KCF, Andrade FA, Bavia L, et al (2019) Chagas Disease: From Discovery to a Worldwide Health Problem. Front Public Heal 7:166. https://doi.org/10.3389/fpubh.2019.00166

14. Mansoldo FRP, Carta F, Angeli A, et al (2020) Chagas Disease: Perspectives on the Past and Present and Challenges in Drug Discovery. Molecules 25:1–14. https://doi.org/10.3390/molecules25225483

15. Jackson Y, Alirol E, Getaz L, et al (2010) Tolerance and safety of nifurtimox in patients with chronic chagas disease. Clin Infect Dis an Off Publ Infect Dis Soc Am 51:e69–75. https://doi.org/10.1086/656917

16. Pinto JP, Machado RSR, Xavier JM, Futschik ME (2014) Targeting molecular networks for drug research. Front. Genet.

17. Schoijet AC, Sternlieb T, Alonso GD (2019) Signal Transduction Pathways as Therapeutic Target for Chagas Disease. Curr Med Chem. https://doi.org/10.2174/0929867326666190620093029

18. Ferri G, Edreira MM (2021) All Roads Lead to Cytosol: Trypanosoma cruzi Multi-Strategic Approach to Invasion

19. McDonough KA, Rodriguez A (2012) The myriad roles of cyclic AMP in microbial pathogens: From signal to sword. Nat Rev Microbiol 10:27–38. https://doi.org/10.1038/nrmicro2688

20. Yan K, Gao L, Cui Y, et al (2016) The cyclic AMP signaling pathway: Exploring targets for successful drug discovery (Review). Mol Med Rep 13:3715–3723. https://doi.org/10.3892/mmr.2016.5005

21. Pavan B, Biondi C, Dalpiaz A (2009) Adenylyl cyclases as innovative therapeutic goals. Drug Discov Today 14:982–991. https://doi.org/10.1016/j.drudis.2009.07.007

22. Pierre S, Eschenhagen T, Geisslinger G, Scholich K (2009) Capturing adenylyl cyclases as potential drug targets. Nat Rev Drug Discov 8:321–335. https://doi.org/10.1038/nrd2827

23. Toya Y, Schwencke C, Ishikawa Y (1998) Forskolin derivatives with increased selectivity for cardiac adenylyl cyclase. J Mol Cell Cardiol 30:97–108. https://doi.org/10.1006/jmcc.1997.0575

24. Alasbahi RH, Melzig MF (2010) Plectranthus barbatus: A review of phytochemistry, ethnobotanical uses and pharmacology part 1. Planta Med 76:653–661. https://doi.org/10.1055/s-0029-1240898

25. Lerner A, Kim DH, Lee R (2000) The cAMP signaling pathway as a therapeutic target in lymphoid malignancies. Leuk Lymphoma 37:39–51. https://doi.org/10.3109/10428190009057627

26. Millar JK, Pickard BS, Mackie S, et al (2005) Genetics: DISC1 and PDE4B are interacting genetic factors in schizoprenia that regulate cAMP signaling. Science (80-) 310:1187–1191.https://doi.org/10.1126/science.1112915

27. Diamant Z, Spina D (2011) PDE4-inhibitors: A novel, targeted therapy for obstructive airways disease. Pulm Pharmacol Ther 24:353–360. https://doi.org/10.1016/j.pupt.2010.12.011

28. Page CP, Spina D (2011) Phosphodiesterase Inhibitors in the Treatment of Inflammatory Diseases. pp 391–414

29. Chiricozzi A, Caposiena D, Garofalo V, et al (2016) A new therapeutic for the treatment of moderate-to-severe plaque psoriasis: Apremilast. Expert Rev Clin Immunol 12:237–249. https://doi.org/10.1586/1744666X.2016.1134319

30. Ahmed A, Boulton S, Shao H, et al (2019) Recent Advances in EPAC-Targeted Therapies: A Biophysical Perspective. Cells 8:. https://doi.org/10.3390/cells8111462

31. Musikant D, Ferri G, Durante IM, et al (2017) Host Epac1 is required for cAMP-mediated invasion by Trypanosoma cruzi. Mol Biochem Parasitol 211:67–70. https://doi.org/10.1016/j.molbiopara.2016.10.003

32. Adesse D, Garzoni LR, Huang H, et al (2008) Trypanosoma cruzi induces changes in cardiac connexin43 expression. Microbes Infect 10:21–28. https://doi.org/10.1016/j.micinf.2007.09.017

33. Calvet CM, Melo TG, Garzoni LR, et al (2012) Current understanding of the Trypanosoma cruzi-cardiomyocyte interaction. Front. Immunol.

34. Rodríguez-Angulo H, Toro-Mendoza J, Marques J, et al (2013) Induction of chagasic-like arrhythmias in the isolated beating hearts of healthy rats perfused with Trypanosoma cruzi-conditioned medium. Brazilian J Med Biol Res 46:58–64. https://doi.org/10.1590/1414-431X20122409

35. Oestreich EA, Wang H, Malik S, et al (2007) Epac-mediated activation of phospholipase Cε plays a critical role in β-adrenergic receptor-dependent enhancement of Ca2+ mobilization in cardiac myocytes. J Biol Chem 282:5488–5495. https://doi.org/10.1074/jbc.M608495200

36. Fazal L, Laudette M, Paula-Gomes S, et al (2017) Multifunctional Mitochondrial Epac1 Controls Myocardial Cell Death. Circ Res 120:645–657. https://doi.org/10.1161/CIRCRESAHA.116.309859

37. Yang H, Xue W, Ding C, et al (2021) Vitexin Mitigates Myocardial Ischemia/Reperfusion Injury in Rats by Regulating Mitochondrial Dysfunction via Epac1-Rap1 Signaling. Oxid Med Cell Longev 2021:. https://doi.org/10.1155/2021/9921982

38. Rastogi S, Pandey MM, Rawat AKS (2016) Traditional herbs: a remedy for cardiovascular disorders. Phytomedicine 23:1082–1089. https://doi.org/10.1016/j.phymed.2015.10.012

39. Popovic-Milenkovic MT, Tomovic MT, Brankovic SR, et al (2014) Antioxidant and anxiolytic activities of Crataegus nigra Wald. et Kit. berries. Acta Pol Pharm - Drug Res 71:279–285

40. Cuevas-Durán RE, Medrano-Rodríguez JC, Sánchez-Aguilar M, et al (2017) Extracts of crataegus oxyacantha and rosmarinus offcinalis attenuate ischemic myocardial damage by decreasing oxidative stress and regulating the production of cardiac vasoactive agents. Int J Mol Sci 18:. https://doi.org/10.3390/ijms18112412

41. Saoudi M, Slama-Ben Salem R Ben, Salem M Ben, et al (2019) Beneficial effects of crataegus oxyacantha extract on neurobehavioral deficits and brain tissue damages induced by an insecticide mixture of deltamethrin and chlorpyrifos in adult wistar rats. Biomed Pharmacother 114:108795. https://doi.org/10.1016/j.biopha.2019.108795

42. Jayalakshmi R, Devaraj SN (2010) Cardioprotective effect of tincture of Crataegus on isoproterenol-induced myocardial infarction in rats. J Pharm Pharmacol 56:921–926. https://doi.org/10.1211/0022357023745

43. Laurent AC, Bisserier M, Lucas A, et al (2015) Exchange protein directly activated by cAMP1 promotes autophagy during cardiomyocyte hypertrophy. Cardiovasc Res 105:55–64. https://doi.org/10.1093/cvr/cvu242

44. Surinkaew S, Aflaki M, Takawale A, et al (2019) Exchange protein activated by cyclic-adenosine monophosphate (Epac) regulates atrial fibroblast function and controls cardiac remodelling. Cardiovasc Res 115:94–106. https://doi.org/10.1093/cvr/cvy173

45. Longo PA, Kavran JM, Kim MS, Leahy DJ (2013) Transient mammalian cell transfection with polyethylenimine (PEI). In: Methods in Enzymology. Academic Press Inc., pp 227–240

46. Che X, Wang X, Zhang J, et al (2016) Vitexin exerts cardioprotective effect on chronic myocardial ischemia/reperfusion injury in rats via inhibiting myocardial apoptosis and lipid peroxidation. Am J Transl Res 8:3319–3328

47. Wu Y, Zhang F, Yang K, et al (2019) SymMap: an integrative database of traditional Chinese medicine enhanced by symptom mapping. Nucleic Acids Res 47:D1110–D1117. https://doi.org/10.1093/nar/gky1021

48. Fang S, Dong L, Liu L, et al (2021) HERB: a high-throughput experiment-and reference-guided database of traditional Chinese medicine. Nucleic Acids Res 49:D1197–D1206. https://doi.org/10.1093/nar/gkaa1063

49. HMPC (2016) Hawthorn leaf and flower

50. Organization WH, Plants WHOC on SM, WHO Consultation on Selected Medicinal Plants (2nd□: 1999□: Ravello-Salerno I, et al WHO monographs on selected medicinal plants

51. American Botanical Council HerbMed Database

52. Elsadig Karar MG, Kuhnert N (2016) UPLC-ESI-Q-TOF-MS/MS Characterization of Phenolics from Crataegus monogyna and Crataegus laevigata (Hawthorn) Leaves, Fruits and their Herbal Derived Drops (Crataegutt Tropfen). J Chem Biol Ther 01: https://doi.org/10.4172/2572-0406.1000102

53. Wang J, Xiong X, Feng B (2013) Effect of crataegus usage in cardiovascular disease prevention: An evidence-based approach. Evidence-based Complement Altern Med 2013:. https://doi.org/10.1155/2013/149363

54. Koch E, Malek FA (2011) Standardized extracts from hawthorn leaves and flowers in the treatment of cardiovascular disorders preclinical and clinical studies. Planta Med 77:1123–1128. https://doi.org/10.1055/s-0030-1270849

55. Parnell E, Palmer TM, Yarwood SJ (2015) The future of EPAC-targeted therapies: agonism versus antagonism. Trends Pharmacol Sci 36:203–214. https://doi.org/10.1016/j.tips.2015.02.003

